# Proteolytic activation of SARS-CoV-2 spike at the S1/S2 boundary: potential role of proteases beyond furin

**DOI:** 10.1101/2020.10.04.325522

**Authors:** Tiffany Tang, Javier A. Jaimes, Miya K. Bidon, Marco R. Straus, Susan Daniel, Gary R. Whittaker

## Abstract

The severe acute respiratory syndrome coronavirus 2 (SARS-CoV-2) uses its spike (S) protein to mediate viral entry into host cells. Cleavage of the S protein at the S1/S2 and/or S2’ site(s) is associated with viral entry, which can occur at either the cell plasma membrane (early pathway) or the endosomal membrane (late pathway), depending on the cell type. Previous studies show that SARS-CoV-2 has a unique insert at the S1/S2 site that can be cleaved by furin, which appears to expand viral tropism to cells with suitable protease and receptor expression. Here, we utilize viral pseudoparticles and protease inhibitors to study the impact of the S1/S2 cleavage on infectivity. Our results demonstrate that S1/S2 pre-cleavage is essential for early pathway entry into Calu-3 cells, a model lung epithelial cell line, but not for late pathway entry into Vero E6 cells, a model cell line. The S1/S2 cleavage was found to be processed by other proteases beyond furin. Using bioinformatic tools, we also analyze the presence of a furin S1/S2 site in related CoVs and offer thoughts on the origin of the insertion of the furin-like cleavage site in SARS-CoV-2.

## Introduction

The 21^st^ century has seen the rise of pathogenic strains of human coronaviruses (CoVs) causing major public health concerns, first with the severe acute respiratory syndrome coronavirus (SARS-CoV) in 2002^1^, then with Middle East respiratory syndrome coronavirus (MERS-CoV) in 2012^2^, and now, with the severe acute respiratory syndrome coronavirus 2 (SARS-CoV-2). SARS-CoV-2 causes the disease syndrome known as COVID-19^3^, now classified as a pandemic with global reach and impact.

CoV host cell entry is mediated by its spike (S) glycoprotein, a large transmembrane protein that decorates the virus particle^4^. The S protein is demarcated into two domains, the S1 and S2 domain, which contains the receptor binding domain and membrane fusion domain, respectively. There are two proteolytic activation events associated with S-mediated membrane fusion^5^. The first is a priming cleavage that occurs at the interface of the S1 and S2 domain (S1/S2) for some coronaviruses, and the second is the obligatory triggering cleavage that occurs within the S2 region (S2’)^5^. The priming cleavage generally converts the S protein into a fusion competent form, by enabling the S protein to better bind receptors or expose hidden cleavage sites^5^. The triggering cleavage initiates a series of conformational changes that enable the S protein to harpoon into the host membrane for membrane fusion^5^.

There are a variety of proteases capable of priming and triggering CoV S proteins. While the priming event is not as well characterized and virus dependent, it has been observed that the CoV S can be triggered by proteases at the plasma membrane or endosomal membrane, enabling entry in what are termed “early” and “late” pathways, respectively^6–8^. SARS-CoV was found to utilize the transmembrane bound protease TMPRSS2 to enter using the early pathway^8–10^. However, TMPRSS2 expression is limited to epithelial cell lines, and in TMPRSS2 negative cell lines, SARS-CoV utilizes endosomal cathepsin L to enter using the late pathway^11^. MERS-CoV also utilizes TMPRSS2 and cathepsin L with one major difference. The MERS-CoV S1/S2 boundary contains an RSVR insert that can be recognized by furin or related proprotein convertases (PC), proteases commonly found in the secretory pathway of most cell lines^7,12^. During the S maturation process, the S protein can be cleaved by furin/PCs^7,12^. Thus, MERS-CoV particles harbor cleaved S1/S2 protein and it was observed that the S1/S2 priming (pre-cleavage) is crucial for MERS-CoV^7^, but not SARS-CoV^13^, to infect via the early pathway route. However, the S1/S2 cleavage is not a requirement for MERS-CoV late pathway infection^7^.

For SARS-CoV-2, early studies showed that the S1/S2 junction contains an insert with two additional basic residues, P-R-R-A (P – proline, R – arginine, A – alanine), that was not present in SARS-CoV or its closest bat ancestor viruses^14,15^. This insert forms a P-R-R-A-R sequence and while it does contain the minimum furin recognition motif, R-X-X-R, it diverges from the preferred R-X-K/R-R motif, because the SARS-CoV-2 sequence has an A in the P3 location instead of an R^16,17^ (**Figure 1**). Intriguingly, the only other known example of this insert on FurinDB^18^, a database of furin substrates, is found in proaerolysin, a bacterial toxin, and was determined to be activated by furin^19^.

**Figure 1:**
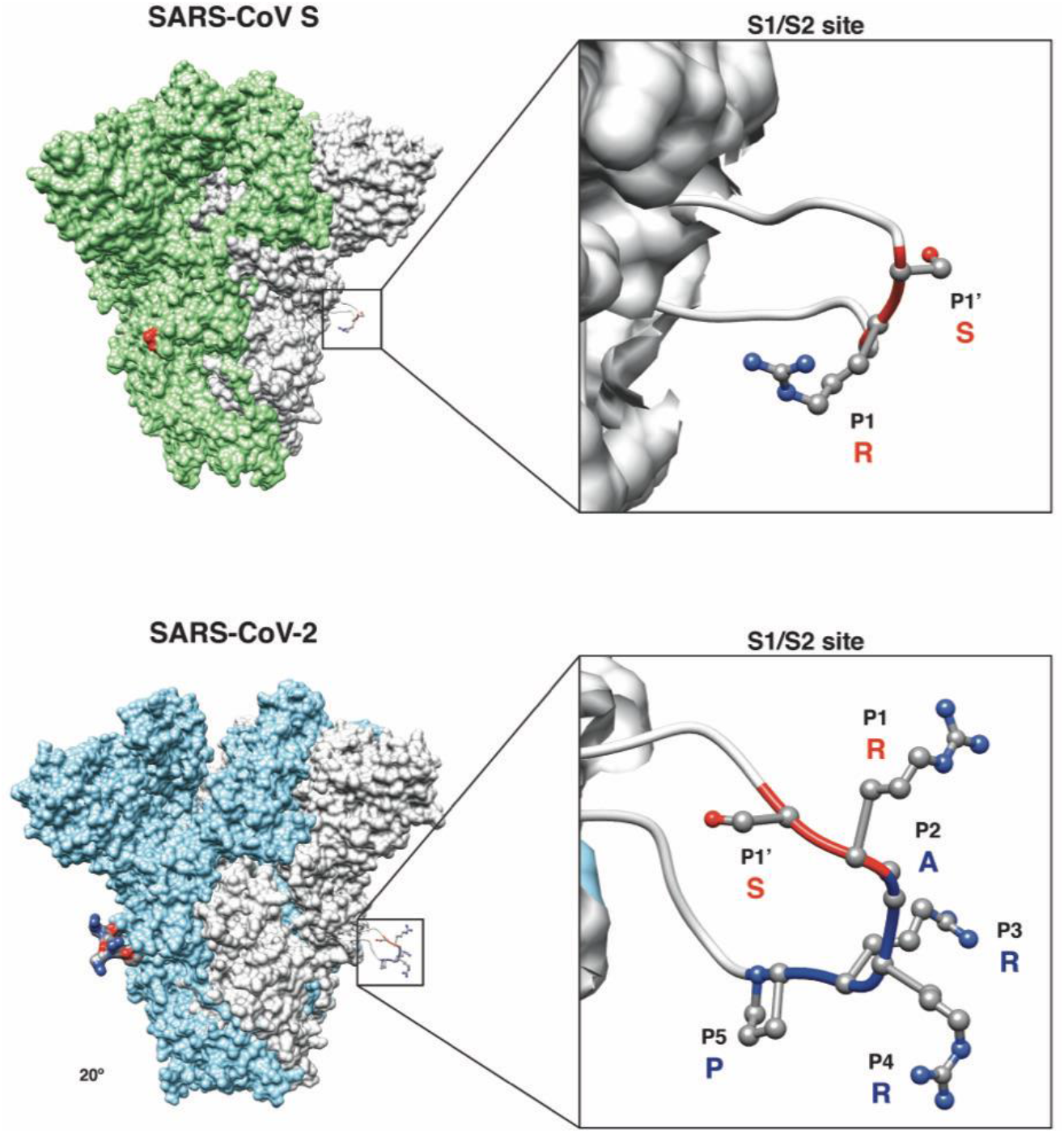
Predicted structural model of the SARS-CoV and SARS-CoV-2 S proteins. (**Inset**) Magnification of the S1/S2 site with conserved R and S residues (red ribbon) and the unique four amino acid insertion P-R-R-A for SARS-CoV-2 (blue ribbon) are shown. The P’s denote the position of that amino acid from the S1/S2 cleavage site, with P1-P5 referring to amino acids before the cleavage site and P1’ referring to amino acids after the cleavage site.

Indeed, early studies suggest that SARS-CoV-2 is also processed by furin since SARS-CoV-2 harbors a cleaved S protein, likely due to furin processing at the S1/S2 site^20–22^. Similar to what was observed for MERS-CoV, this S1/S2 cleavage was determined to be a prerequisite for subsequent TMPRSS2 activation at the S2’ site for infection of respiratory cell lines, such as Calu-3^23,24^. Furthermore, like MERS-CoV, SARS-CoV-2 can also utilize cathepsin L in the endosomal pathway in TMPRSS2-negative cell lines, such as Vero E6^22^.

In this study, we sought to better characterize the SARS-CoV-2 S1/S2 cleavage site and raise some intriguing possibilities for how the site might have emerged. Using pseudoparticles and protease inhibitors, we investigated the impact of S1/S2 cleavage for successful infection of cells via the early and late pathway. Our data shows that the S1/S2 cleavage is essential for subsequent S2’ activation via TMPRSS2 for entry in Calu-3 cells, but not for S2’ activation via cathepsin L for entry in Vero E6 cells, and suggests that furin may not be the only protease responsible for the S1/S2 cleavage event.

## Results and Discussion

### MLV pseudoparticles as a system to study SARS-CoV-2 entry

To assess the functional importance of the S1/S2 site for SARS-CoV-2 entry, we utilized viral pseudoparticles. These particles consist of a murine leukemia virus (MLV) core and are decorated with the viral envelope protein to accurately recapitulate the entry steps of their native counterpart^25^. These particles also contain a luciferase reporter that integrates into the host cell genome upon successful infection and drives the cells to produce luciferase, which is quantifiable. We and others have used MLV-based pseudoparticles widely to study CoV entry pathways^12,21,26^. Using the HEK293T cell line, MLV pseudoparticles containing the wild type SARS-CoV-2 S protein (SARS-CoV-2pp), or the SARS-CoV S protein (SARS-CoVpp) were generated alongside positive control particles containing the vesicular stomatitis virus G protein (VSVpp) or negative control particles lacking envelope proteins (Δenvpp).

Since coronaviruses can enter via the “early” or “late” pathway, we chose to infect cell lines representative of each pathway, as the entry mechanism can be highly cell-type dependent^27^. We utilized the Calu-3 (early) and the Vero E6 (late) cell lines for these studies, which activate the SARS-CoV-2 S2’ using TMPRSS2 and cathepsin L, respectively. As expected, VSVpp (positive control) infected Vero E6 and Calu-3 cells with several orders of magnitude higher luciferase units than the values reported with Δenvpp infection (negative control) (**Figure 2A, 2B**). This confirms that the envelope protein is driving infection, and not the particle itself. In the case of SARS-CoVpp and SARS-CoV-2pp, both particles are infectious in both cell lines as they drive luciferase production several orders of magnitude higher than Δenvpp. (**Figure 2A, 2B**), with SARS-CoVpp more infectious than SARS-CoV-2pp.

**Figure 2:**
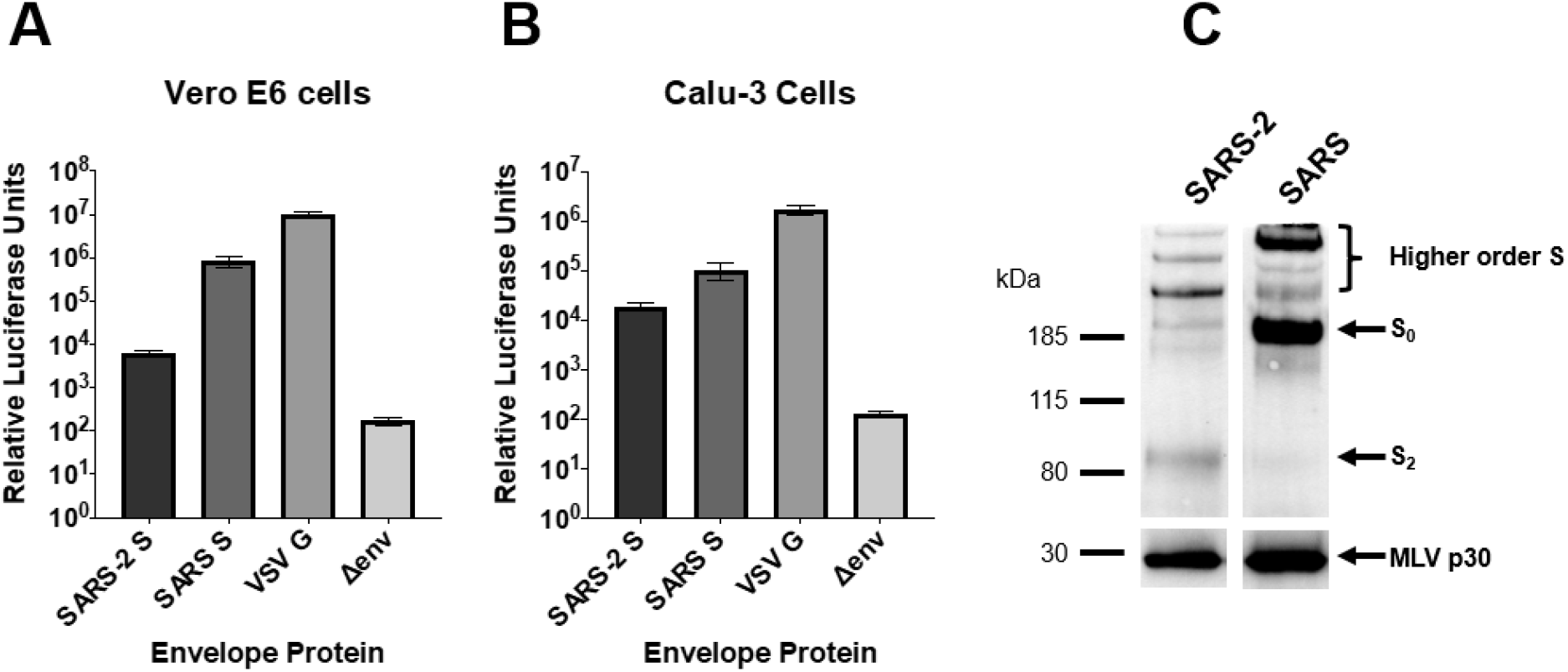
Characterization of MLVpp system. (**A and B**) MLVpp infectivity in Vero E6 and Calu-3 cells. Cells were infected with MLVpps exhibiting the SARS-CoV-2 S, SARS-CoV S, VSV G or no envelope protein and assessed for luciferase activity. Error bars represent the standard error measurements of three biological replicates (n=3). (**C**) Western blot analysis of SARS-CoV-2pp and SARS-CoVpp S content. S was detected using a rabbit antibody against the SARS-CoV-2 S2 region and cross reacts against the SARS-CoV S. MLV content was detected using a mouse antibody against MLV p30. Image cropped from a singular western blot.

The pseudoparticles were also probed for their S content via western blot. For the SARS-CoV-2pp, we detected a band at 85 kDa and for SARS-CoVpp, a strong band at 185 kDa (**Figure 2C**). The different bands observed between SARS-CoV-2pp and SARS-CoVpp is likely because SARS-CoV-2 S is cleaved at the S1/S2 location by furin/PCs during biosynthesis, while SARS-CoV S cannot be cleaved. Thus, the 85 kDa band corresponds to the S2 segment (S_2_) of the SARS-CoV-2 S, as the primary antibody recognizes the S2 region, while the 185 kDa band corresponds to the uncleaved (S_0_) SARS-CoV S. We note that the both SARS-CoVpp and SARS-CoV-2pp exhibit bands at >185 kDa that likely corresponding to dimeric and trimeric forms of S. These oligomeric forms of S have been reported in other SARS-CoV-2pp western blots^22,28^. Since SARS-CoVpp harbor full-length S, their oligomeric S bands are of higher molecular weight than the SARS-CoV-2pp oligomeric S bands, which harbor cleaved S. Furthermore, the higher order bands may be more apparent due to the use of a polyclonal antibody to detect both the SARS-CoV and SARS-CoV-2 S.

### Use of the dec-RVKR-CMK protease inhibitor to produce SARS-2pp with uncleaved S

To examine the functional role of S priming by furin/PC proteases, we needed to produce SARS-CoV-2pp expressing uncleaved S protein. This can be accomplished by treating producer cells with an appropriate protease inhibitor to prevent S1/S2 cleavage during biogenesis. We chose dec-RVKR-CMK because it has been shown to inhibit furin/PCs, preventing S1/S2 cleavage^23,26,29^. We have previously showed that addition of dec-RVKR-CMK to producer cells generates MLV pseudotyped particles that harbor full-length MERS-CoV S protein (MERS-CoVpp)^26^ and we observe a similar trend with SARS-CoV-2pp generated from cells treated with dec-RVKR-CMK (**Figure 3C,** lanes 1 and 2), exhibiting full-length uncleaved S_0_ and weak cleaved S_2_ bands. As mentioned, the additional bands at >185 kDa likely corresponding to dimeric and trimeric S, as previously reported in other western blots probing for SARS-CoV-2pp S^22,28^, with the higher order bands from particles produced in cells treated with dec-RVKR-CMK observed at higher molecular weights.

**Figure 3:**
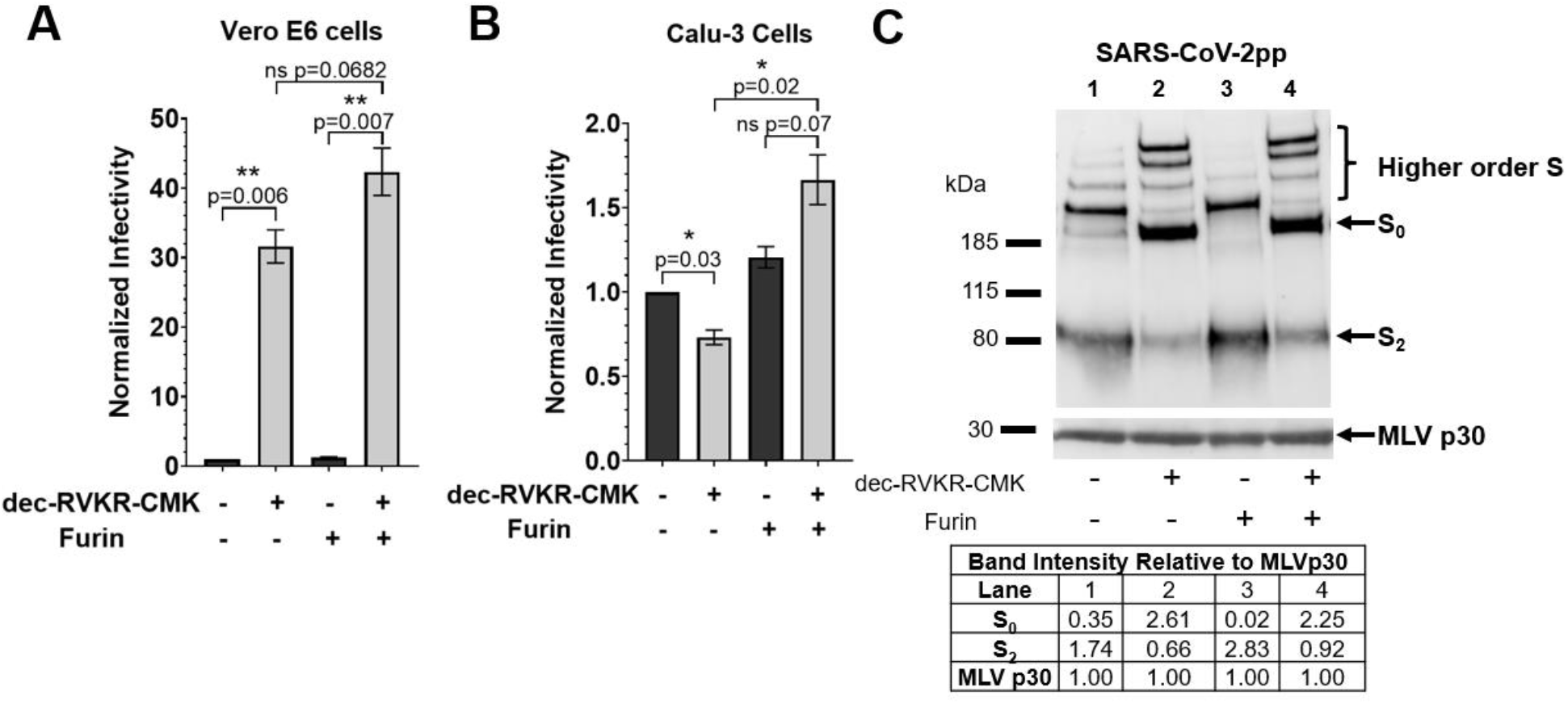
Impact of dec-RVKR-CMK on MLVpps. + dec-RVKR-CMK refers to MLVpp produced in HEK293T treated with 75 μM of dec-RVKR-CMK at the time of transfection. + Furin refers to particles treated with 6 U of recombinant furin for 3 hours at 37°C. (**A and B**) +/− dec-RVKR-CMK and +/− Furin SARS-CoV-2pp were used to infect Vero E6 and Calu-3 cells. Infectivity was normalized to the – dec-RVKR-CMK – furin condition in each cell line. Note the change in y-axis scale between A and B. Error bars represent the standard error measurements of three biological replicates (n=3). Statistical analysis was performed using an unpaired student’s t test. Not shown: there is a ns difference between the – dec-RVKR-CMK +/− furin conditions (black bars). *, P ≤ 0.05, **, P ≤ 0.01, ns, non-significant, P > 0.05. (**C**) Western blot analysis of SARS-CoV-2pp produced in +/− dec-RVKR-CMK and treated with +/− furin for S protein content to complement the infection conditions. S was detected using a rabbit antibody against the SARS-CoV-2 S2 region. MLV content was detected using a mouse antibody against MLV p30. Band intensities were normalized to the MLV p30 band intensity.

The dec-RVKR-CMK-treated (uncleaved) particles were used to transduce cells to observe the impact of S1/S2 cleavage for subsequent S2’ activation via early acting TMPRSS2 in Calu-3 cells or late acting cathepsin L in Vero E6 cells. In Vero E6 cells, the uncleaved SARS-CoV-2pp were 30-fold more infectious than their cleaved counterparts, suggesting that uncleaved particles result in more infectious particles, and that the S1/S2 pre-cleavage is not required or hinders infection via the late pathway (**Figure 3A**). However, in Calu-3 cells, the opposite trend was observed; the uncleaved particles were significantly less infectious than their cleaved counterparts (**Figure 3B**), suggesting that S1/S2 pre-cleavage is essential^23,24^ for early entry. As a control, we also generated SARS-CoVpp, which lacks the furin/PC S1/S2 site, in the presence or absence of dec-RVKR-CMK and noted no significant impact of dec-RVKR-CMK on SARS-CoVpp infectivity in both Vero E6 and Calu-3 cells (**Figure S1A and S1B**). This affirms that the dec-RVKR-CMK impacts on infectivity is due to blocking S1/S2 cleavage.

Thus, we were interested to see if we could restore the infection of the uncleaved particles by treating them with exogeneous furin, which has been previously shown to process purified SARS-CoV-2 S protein^30^. In Calu-3, we observed a 2-fold increase in infection when uncleaved particles were treated with exogeneous furin, but with cleaved particles, we did not observe any statistically significant increase with furin treatment (**Figure 3B**). In Vero E6 cells, for both types of particles, we did not observe a statistically significant increase in infection with furin treatment. (**Figure 3A**).

Since furin treatment only modestly increased infectivity in all cases observed, we wanted to visualize how efficiently furin was processing the SARS-CoV-2 S via western blot. For cleaved particles, exogeneous furin processed the faint full-length S_0_ band into S_2_ (**Figure 3C**, lanes 1 and 3). This trend was confirmed upon quantifying the intensities of the S_0_ and S_2_ bands relative to the intensity of the MLV p30 band; furin treatment reduced the relative S_0_ intensity from 0.35 to 0.02, resulting in an increase in relative S_2_ intensity from 1.74 to 2.83. For uncleaved particles, furin treatment had a more modest impact, as the relative intensity of the S_0_ bands was reduced from 2.61 to 2.25, resulting in a small increase in relative S_2_ intensity from 0.66 to 0.92 (**Figure 3C**, lanes 2 and 4).

The western blot bands also reveal that the relative intensity of the uncleaved SARS-CoV-2pp S_0_ bands are greater than the relative intensity of the cleaved SARS-CoV-2pp S_2_ bands and this trend was previously observed with MERS-CoVpp S^26^ (**Figure 3C**, lanes 1 and 2), but not with SARS-CoV (**Figure S1C**). This likely suggests that the increased infectivity we observed with the uncleaved SARS-CoV-2pp in Vero E6 cells could be due to particles harboring a greater amount of S protein. A possible explanation why uncleaved particles could harbor more S is because uncleaved S is more stable than cleaved S, as S1/S2 cleavage reduces S stability^30^. Thus, cleavage may encourage greater S degradation, reducing the amount of S available for incorporation into pseudoparticles. However, we did not observe increased infectivity with uncleaved SARS-CoV-2pp in Calu-3 cells because of the S1/S2 cleavage requirement and increased S content cannot compensate for cleavage.

Due to the limited ability of exogenous furin to rescue and cleave SARS-CoV-2 S, we re-evaluated the role of furin in processing the S1/S2 site. Since dec-RVKR-CMK can also inhibit a number of furin related PCs, the effects that have been observed with dec-RVKR-CMK on particles could have resulted from inhibition of other PC proteases and not necessarily furin. Therefore, we produced particles in the presence of alpha1-PDX, a more selective furin inhibitor^31^ than dec-RVKR-CMK. Infectivity results of these particles in Vero E6 cells show that the alpha1-PDX furin specific inhibitor failed to recapitulate the high level of enhancement provided by dec-RVKR-CMK at all tested inhibitor concentrations (**Figure 4A**). This indicates that dec-RVKR-CMK may have also been acting on non-furin proteases that can also process the S1/S2 site and furin itself may not be the only protease processing the S1/S2 site, as is generally assumed.

**Figure 4:**
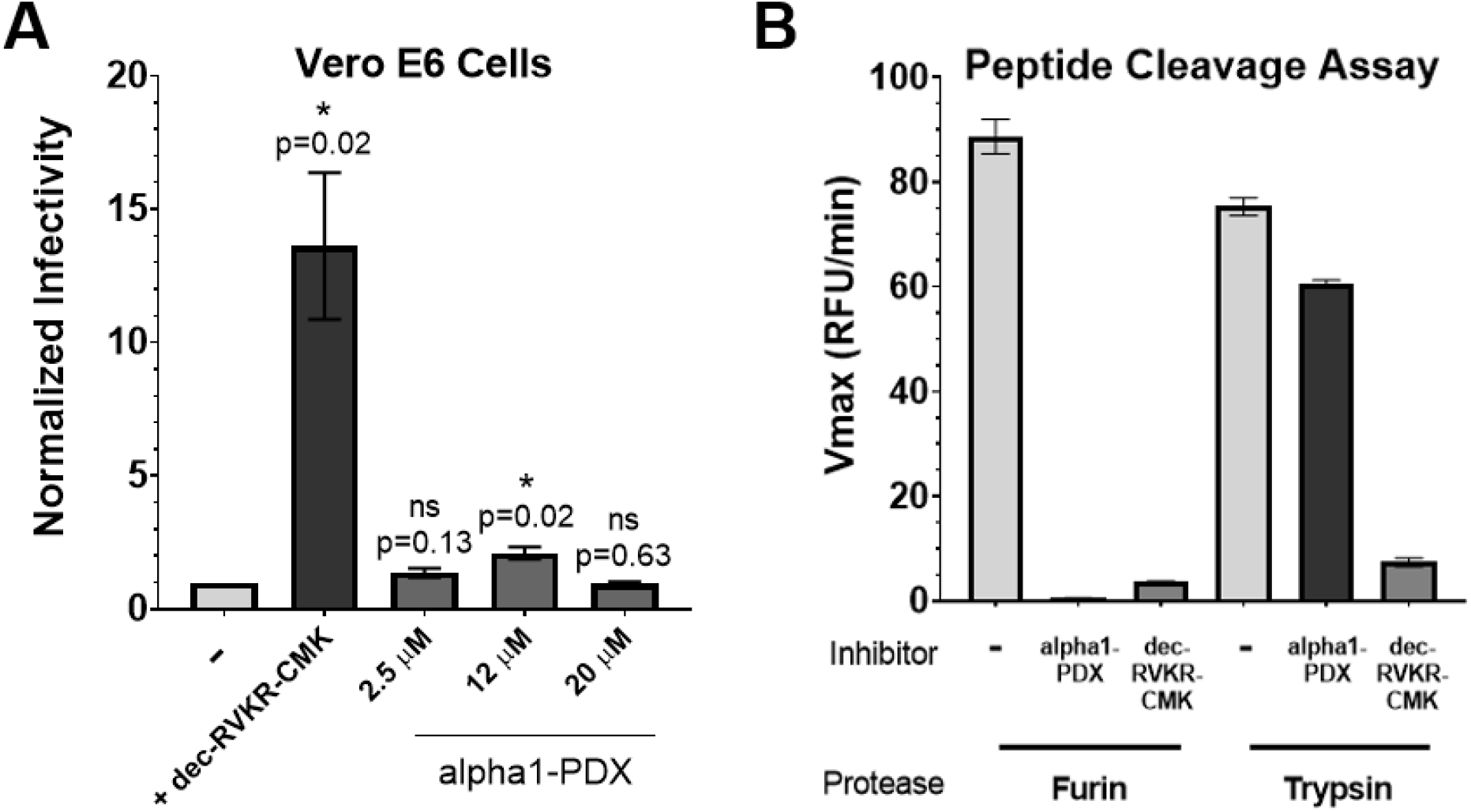
Use of specific furin (alpha1-PDX) and broad furin (dec-RVKR-CMK) inhibitors to investigate furin specificity. **(A)** Infectivity of particles produced in the presence of broad (dec-RVKR-CMK) and specific (alpha1-PDX) furin inhibitors. Inhibitor was applied at the indicated concentration to producer HEK293T cells. Particles were used to infect Vero E6 cells. Infectivity was normalized to the – condition. Error bars represent the standard error measurements of four biological replicates (n=4). Statistical analysis was performed using an unpaired student’s t test compared to - condition. *, P ≤0.05, ns, non-significant, P > 0.05. **(B)** Proteolytic cleavage assay of SARS-CoV-2 S1/S2 peptides to confirm alpha1-PDX and dec-RVKR-CMK inhibition specificity. Alpha1-PDX (2 μM), dec-RVKR-CMK (75 μM) or no inhibitor was incubated with either furin or trypsin recombinant protease and a fluorogenic peptides mimicking the SARS-CoV-2 S1/S2 site (TNSPRRARSVA). Results represent averages and error bars represent the standard error measurement of three biological replicates (n=3).

As a control, the activity of the alpha1-PDX was confirmed using a proteolytic cleavage assay, in which proteolytic cleavage of fluorogenic peptides results in a fluorescence increase used to determine the velocity of the cleavage reaction (Vmax). We have previously used this assay to observe furin cleavage of fluorogenic peptides mimicking the SARS-CoV-2 S1/S2 site (TNSPRRARSVA)^32^. Addition of the alpha1-PDX or dec-RVKR-CMK inhibitor to a peptide reaction including furin resulted in almost complete inhibition of activity, as very limited cleavage was detected. Interestingly, addition of the alpha1-PDX or dec-RVKR-CMK to a reaction including trypsin gave different results. While alpha1-PDX did modestly inhibited trypsin-mediated cleavage, dec-RVKR-CMK exhibited almost complete inhibition of trypsin activity. Thus, the results of this peptide cleavage assay confirm that alpha1-PDX is active on furin-like substrates and is a much more specific inhibitor compared to dec-RVKR-CMK (**Figure 4B**).

Our published results demonstrated that the PC protease PC1 can process SARS-CoV-2 S1/S2^32^, which opens up the possibility that non-furin proteases within the secretory pathway in producer cells can process the SARS-CoV-2 “furin-like” S1/S2 site, as originally suggested by Coutard et al^33^. In the studies reported here, we utilize HEK293T cells to produce the pseudoparticles, which may express a different repertoire of proteases than those in clinically relevant cells. In the future, it will be important to investigate the impact of S1/S2 processing by non-furin PCs in cells infected with SARS-CoV-2.

### Furin-cleavage predictions of the SARS-CoV-2 S1/S2 site

To gain further insights on furin processing of the S1/S2 sites, we utilized the PiTou^34^ and ProP^35^ cleavage prediction tools to analyze the likelihood that the S1/S2 site is processed by furin. ProP predicts furin cleavage sites based on networks derived from experimental data, whereas PiTou uses a combination of a hidden Markov model and biological knowledge-based cumulative probability score functions to characterize a 20 amino acid motif from P14 to P6’ that reflects the binding strength and accessibility of the motif in the furin binding pocket. Thus, the PiTou algorithm has been reported to be more sensitive and specific than ProP for predicting furin cleavage^34^. Both algorithms agree with each other in predicting furin cleavage, with positive scores (PiTou) and scores above 0.5 (ProP) indicating furin cleavage, strengthening the predictions (**Figure 5**), but since the PiTou algorithm is more sensitive and specific, we decided to focus on the PiTou values for further analysis.

**Figure 5.**
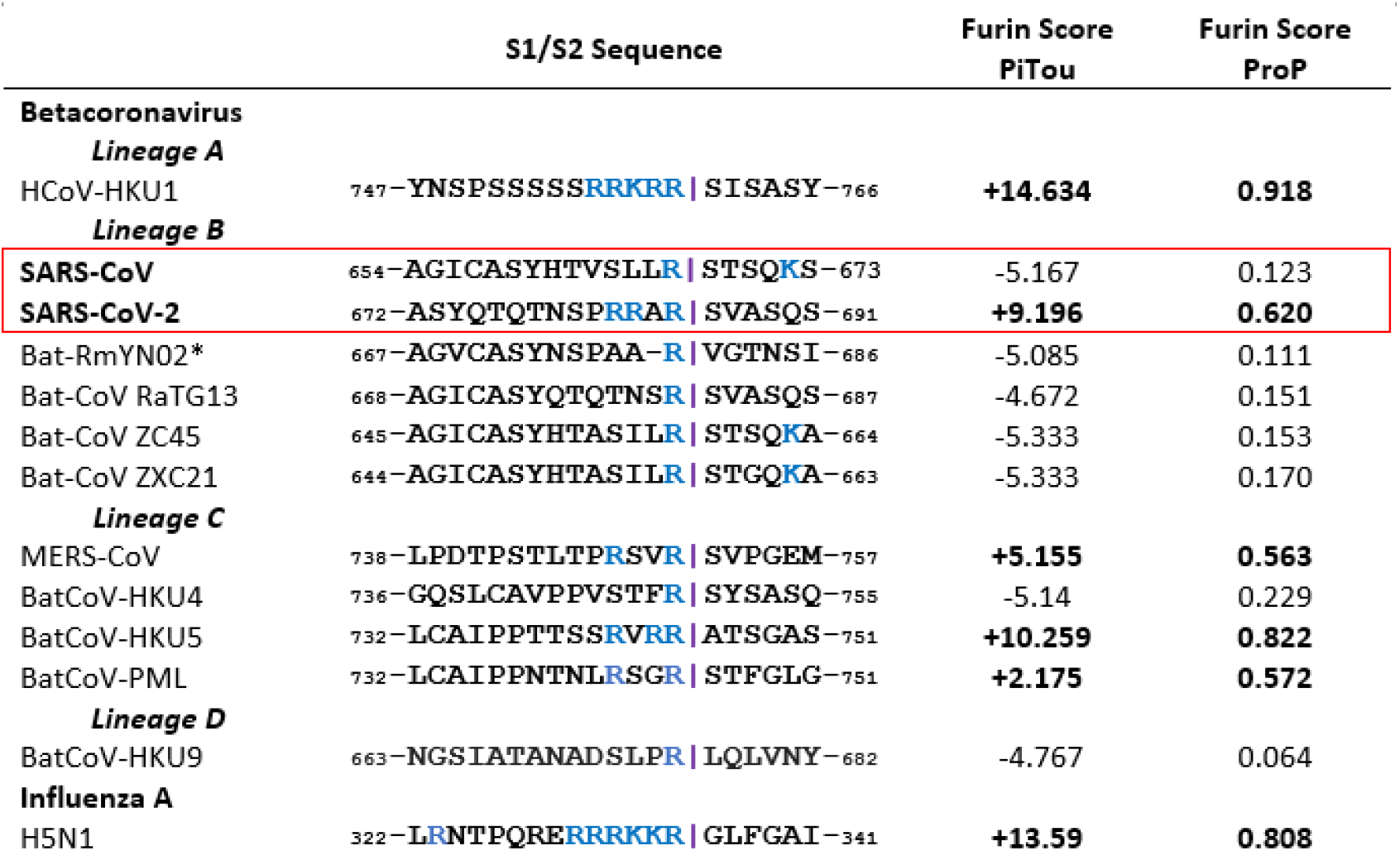
Furin cleavage score analysis of CoV S1/S2 cleavage sites. CoV S sequences were analyzed using the ProP^35^ 1.0 and PiTou^34^ 3.0 furin prediction algorithm. Bold scores indicate the S1/S2 sequence is predicted to be cleaved by furin. Purple lines denote the position of the predicted S1/S2 cleavage site. Basic arginine (R) and lysine (K) residues are highlighted in blue. *For Bat-RmYN02, sequence number was determined from S alignment with SARS-CoV-2 S using Geneious. Sequences corresponding to the S1/S2 region of HCoV-HKU1 (AAT98580.1), SARS-CoV (AAT74874.1), SARS-CoV-2 (QHD43416.1), Bat-CoVRaTG13 (QHR63300.2), Bat-SL-CoVZC45 (AVP78031.1), Bat-SL-CoVZXC21 (AVP78042.1), MERS-CoV (AFS88936.1), BatCoV-HKU4 (YP_001039953.1), BatCoV-HKU5 (YP_001039962.1), BatCoV-PML (KC869678), Bat-CoV-HKU9 (YP_001039971), and Influenza A/Chicken/Hong Kong/822.1/01/H5N1 (AF509026.2) were obtained from GenBank. Sequences corresponding to the S1/S2 region of RmYN02 (EPI_ISL_412977) was obtained from GISAID.

The PiTou algorithm predicts that the SARS-CoV-2 S1/S2 site (PiTou score: 9.2) can be cleaved by furin, whereas other lineage B CoVs that have been proposed as SARS-CoV-2 precursors, such as SARS-CoV, RaTG13, ZC45, ZXC21 and RmYN02, are predicted not to be cleaved by furin. Notably, the PiTou scores of traditionally accepted furin cleavage sites, such as those found in influenza H5N1 (PiTou score: 13.6) and HCoV-HKU1 (PiTou score: 14.6), are much higher than for SARS-CoV-2, suggesting that while SARS-CoV-2 S1/S2 site has increased furin recognition when compared to other lineage B-betaCoVs, the site is not optimal for furin cleavage, in alignment with the experimental data shown here suggesting poor furin specific cleavage.

The modest SARS-CoV-2 PiTou score raises question about the origin and evolution of the S1/S2 site of the virus. It is generally believed that the SARS-CoV-2 S1/S2 site is an insertion of “polybasic” residues^36^, as occurs for avian influenza virus, but as the identity of the SARS-CoV-2 ancestor virus still remains to be discovered, it is important to explore alternative explanations. It has recently been proposed that unsampled SARS-like CoV lineages(s) may be circulating in bats, which are important for the evolutionary origins of SARS-CoV-2^37^. Of note, a “breakpoint sequence hypothesis” has been proposed for the introduction of a furin-like cleavage site at S1/S2^38^, an idea further elucidated by Lytras et al^39^ in the context of the origin of SARS-like viruses. We consider that SARS-CoV-2 could have originated via recombination with a currently unknown ancestor bat virus with a robust furin cleavage site (i.e. PiTou score approx. 13-15), but one which cannot bind to a human receptor—and that in order to gain the ability to infect human cells, the furin cleavage site may have been down-regulated.

These suggested parallels are informed by betaCoV lineage C in which MERS-CoV has a modest furin score (PiTou score: 5.2), but its closest known ancestor bat viruses, BatCoV-HKU4 (PiTou score: <0), BatCoV-HKU5 (PiTou score: 10.3), and BatCoV-PML (PiTou score: 2.175) exhibit a wide range of furin scores. Of note is that the virus with the highest furin score, BatCoV-HKU5 (PiTou score: 10.3), cannot bind the MERS-CoV hDPP4 receptor, whereas BatCoV-HKU4 (PiTou score: <0) can^40^, so the MERS-CoV S1/S2 site can be seen as a down-regulation from BatCoV-HKU5 to bind hDPP4.

Overall, our results support observations^23,24^ that the role of this “gained” SARS-CoV-2 S1/S2 site is to expand viral tropism to respiratory cells, likely in the upper airways, allowing efficient transmission. Cleaved S1/S2 site is crucial for the SARS-CoV-2 S to be subsequently cleaved at the S2’ location by TMPRSS2 for early entry in respiratory cells. Without the S1/S2 pre-cleavage, we predict that SARS-CoV-2 would be endocytosed and due to low cathepsin L expression in respiratory endosomes^7^, SARS-CoV-2 would not efficiently infect via this route. If SARS-CoV-2 infects TMPRSS2 negative cells, it can utilize endosomal cathepsin L, an ubiquitous protease generally found throughout mammalian cells^41^, to activate the S protein^22^—though at undetermined sites. However, for cathepsin L activation, S1/S2 pre-cleavage is not required, and our results indicate that preventing this cleavage increases infectivity of SARS-CoV-2, at least in Vero E6 cells. This may be connected to recent work showing that S1/S2 pre-cleavage reduces the thermal stability of SARS-CoV-2 S^30^ and by preventing this cleavage, there is more S available for incorporation into pseudoparticles. Interestingly, the S1/S2 site activation appears to also have a role in the immune response against SARS-CoV-2, as it has been shown that SARS-CoV-2 S with a deleted S1/S2 loop can provide better protective immune response than with the S1/S2 loop^42^.

The role of furin in activating the S1/S2 site was also investigated. As protease inhibitors commonly employed in cell entry studies, such as dec-RVKR-CMK, can also inhibit other proteases in addition to furin^29^, it is difficult to ascertain the impact of individual proteases. While furin likely processes the SARS-CoV-2 site, our data suggests that other PCs are also involved, since western blots show poor processing of uncleaved S upon addition of purified furin and the use of a highly selective furin inhibitor (alpha1-PDX) has a modest impact on infectivity. All in all, as we consider protease-based inhibitors in treating COVID-19, especially those targeting furin^43,44^, it is important to thoroughly evaluate the role of these proteases to be certain that the relevant proteases are targeted.

## Methods

### Predicted structural modeling

S protein models were built using UCSF Chimera (v.1.14, University of California) through modeler homology tool of the Modeller extension (v.9.25, University of California) as described^15^. SARS-CoV and SARS-CoV-2 S models were built based on the SARS-CoV S structure (PDB No. 5X58).

### Cells, plasmids, and reagents

HEK293T (accession no: CRL-11268, ATCC), Vero E6 (accession no: CRL-1586, ATCC), and Calu-3 (accession no: HTB-55) cells were maintained at 37°C and 5% CO2 incubator and cultivated in Dulbecco’s Modified Eagle Medium (Cellgro) supplemented with 10% Hyclone FetalClone II (GE) and 10 mM HEPES (Cellgro) (DMEMc). Cells were passaged using Dulbecco’s Phosphate-Buffered Saline (DPBS) and Trypsin EDTA 1x (Cellgro). The pCMV-MLV-gagpol murine leukemia virus (MLV) packaging construct, the pTG-luc luciferase reporter, pCAGGS/VSV-G, pCAGGS, and pcDNA/SARS-S plasmids are as previously described^25^. The pcDNA/SARS2-S was a generous gift from Veesler lab^21^.

Protease inhibitor decanoyl-RVKR-CMK (dec-RVKR-CMK) was purchased from Tocris and resuspended in sterile water to 10 mM. Recombinant furin was purchased from New England Biolabs. Recombinant L-1-Tosylamide-2-phenylethyl chloromethyl ketone (TPCK)-treated trypsin was obtained from Sigma-Aldrich. Human alpha-1 PDX (alpha1-PDX) recombinant protein was purchased from ThermoFisher Scientific.

The SARS-CoV-2 S1/S2 fluorogenic peptide with the sequence TNSPRRARSVA, sandwiched between the (7-methoxycoumarin-4-yl)acetyl/2,4-dinitrophenyl (MCA/DNP) FRET pair, was synthesized by Biomatik.

### Furin prediction calculations

<u>Prop</u>: CoV sequences were inputted into the ProP 1.0 Server hosted here: cbs.dtu.dk/services/ProP/ <u>PiTou</u>: CoV sequences were analyzed using the PiTou V3 software freely available via request here^34^.

### Pseudoparticle production

Pseudotyped virus were produced from published protocols with minor modifications^25^. Briefly, HEK293T cells were seeded to 30% confluency. Cells were transfected with pTG-luc (600 ng), pCMV-MLV-gagpol (800 ng), and the respective viral envelope protein (600 ng) using polyethylenimine for 48 hours. Supernatants were then harvested, centrifuged, clarified, and stored in - 80°C aliquots.

For + dec-RVKR-CMK particles, 7.5 μL of dec-RVKR-CMK was added to cells immediately after transfection and boosted with an additional 7.5 μL 24 hours later (final concentration: 75 μM).

For alpha1-PDX particles, indicated concentrations were added to cells immediately after transfection, as recommended by the manufacturer.

### Pseudoparticle assays

Infection assays were as previously described with minor modifications^25^ in triplicate. Briefly, target cells were seeded to confluency. Cells were washed with DPBS, infected with particles on a rocker in an incubator with rocker for 1.5 hours. Cells were supplemented with DMEMc and incubated for 72 hours. Cells were lysed and luciferase activity was assessed using the Luciferase Assay System and the Glomax 20/20 luminometer (Promega). Infectivity values were analyzed and plotted using Prism 8.

### Exogeneous protease treatment

1 mL pseudoparticles were pelleted using a TLA-55 rotor with an Optima-MAX-E ultracentrifuge (Beckman Coulter) for 2 hours at 42,000 rpm at 4°C. Particles were resuspended in 30 μL DPBS buffer supplemented with 20 mM HEPES, 0.2 mM CaCl2, 0.2 mM β-mercaptoethanol (at pH 7.0). Particles were treated with 6 U recombinant furin for 3 hours at 37°C.

### Western blot analysis of pseudoparticles

3 mL pseudoparticles were pelleted and resuspended as described above, but in 20 μL. Sodium dodecyl sulfate (SDS) loading buffer and DTT were added to samples and heated at 65°C for 20 minutes. Samples were separated on NuPAGE Bis-Tris gel (Invitrogen) and transferred on polyvinylidene difluoride membranes (GE). SARS-CoV-2 and SARS-CoV S was detected using a rabbit polyclonal antibody that recognizes the S2 domain (Cat: 40590-T62, Sinobiological) and an AlexaFluor 488 goat anti-rabbit secondary antibody. MLV protein was detected using a mouse monoclonal antibody that recognizes the MLV p30 protein (Cat: ab130757, abcam) and an AlexaFluor 488 goat antimouse secondary antibody. Bands were detected using the ChemiDoc Imaging software (Bio-Rad). S_0_ and S_2_ band intensity was calculated using the analysis tools on Biorad Image Lab 6.1 software and normalized to the MLV p30 band intensity.

### Fluorogenic peptide assay

Peptide assays were as previously described with minor modifications^32^ in triplicates. Each reaction was performed in a 100 μL volume consisting of buffer, protease, inhibitor, and SARS-CoV-2 S1/S2 fluorogenic peptide in an opaque 96-well plate. For furin catalyzed reactions, 1 U/well recombinant furin was diluted in buffer consisting of 20 mM HEPES, 0.2 mM CaCl_2_, 0.2 mM β-mercaptoethanol. For trypsin catalyzed reactions, 0.8 nM/well TPCK trypsin was diluted in PBS buffer. The alpha1-PDX inhibitor or dec-RVKR-CMK inhibitor were added at a concentration of 2 μM or 75 μM, respectively. Peptide was lastly added at a concentration of 50 μM.

Fluorescence emission was measured every minute for 60 minutes using a SpectraMax fluorometer (Molecular Devices) at 30°C using an excitation wavelength of 330 nm and an emission wavelength of 390 n. Vmax was calculated by fitting the linear rise in fluorescence to the equation of a line.

## Acknowledgments

This work was funded by the National Institute of Health research grant R01AI35270 and Fast Grant, Mercatus Center. TT acknowledges support by the National Science Foundation Graduate Research Fellowship Program under Grant No. DGE-1650441 and the Samuel C. Fleming Family Graduate Fellowship. We would especially like to thank Jean Millet for providing important insight on analyzing PiTou values and calculations for BatCoV-HKU9. We would like to thank Hector Aguilar-Carreno for helpful input and thank members of the Daniel and Whittaker groups as well as the Eliezer and Weinstein groups at Weill Cornell for helpful discussions. SD acknowledges funding for this project, sponsored by the Defense Advanced Research Projects Agency (DARPA) Army Research Office and accomplished under Cooperative Agreement Number W911NF-18-2-0152. The views and conclusions contained in this document are those of the authors and should not be interpreted as representing the official policies, either expressed or implied, of DARPA or the Army Research Office or the U.S. Government. The U.S. Government is authorized to reproduce and distribute reprints for Government purposes notwithstanding any copyright notation herein.

## Supporting Information

Supporting information available: Impact of dec-RVKR-CMK inhibitor on SARS-CoVpp production, which lacks the furin S1/S2 site (**Figure S1**).

